# Consolidated Calcium kinetic rates in a Caucasian population as a function of age and sex

**DOI:** 10.1101/2024.06.18.599503

**Authors:** Niklas Hartung, Steven A Abrams, Wilhelm Huisinga, Karin Weisser

## Abstract

Calcium plays an important role in bone physiology and its kinetics change over lifetime. The analysis of calcium deposition and resorption through stable isotope techniques has guided recommendations on nutritional uptake for overall health. In addition, calcium kinetics have great relevance for toxicokinetic studies of bone-seeking elements (e.g, aluminium and lead) since these elements use common uptake and release pathways. While the impact of many factors on calcium kinetics have been investigated individually, a consolidated age- and sex-dependent kinetic description amenable for toxicokinetic modeling, however, is still lacking. Motivated by this need, we systematically reviewed the existing literature on calcium kinetics and assembled a large and consistent dataset. Then, building on the work of O’Flaherty in the 1990s, we formulated age- and sex-dependent functions describing calcium deposition, resorption, net retention, and mass. This description represents the current knowledge on calcium kinetics in a reference individual of Caucasians as most data was from this population.

## 1. INTRODUCTION

Calcium deposition to and release from bone play an important role in bone physiology. Between birth and adulthood, these kinetic processes change considerably, peaking during puberty [1]. The rate of change of total calcium mass, called calcium net retention rate (sometimes also calcium accretion rate), results as the net balance between deposition and release rates.

Since the advent of stable isotope methods for quantification of calcium kinetics in humans in the 1980s [2], these processes have been analyzed extensively in the literature, including the impact of sex, ethnicity, age, and various diseases on calcium kinetics [1, 2]. Also, dual-energy X-ray absorptiometry and neutron activation analysis have rendered feasible the in vivo quantification of whole-body calcium [3], and reference values for whole-body calcium have been reported for males and females of different ages and ethnicities [4].

Besides the importance of calcium absorption and kinetics for guiding recommendations on nutritional calcium uptake [5], calcium kinetic processes are also relevant for toxicokinetic evaluations of bone-seeking metals like lead or aluminium. Indeed, several authors have formulated toxicokinetic models on the assumption that bone distribution of bone-seeking metals parallels calcium kinetics [6, 7, 8, 9, 10].

Several efforts to consolidate calcium kinetic parameters have been reported in the literature. For selected age groups (preterm, newborn, pre-/post-pubertal, adult), sex-independent calcium deposition rates have been proposed [1]. The last effort to derive age-continuous functions from the diverse literature on calcium kinetics dates back more than 25 years [11]. While a large body of newer literature on the topic (most calcium kinetic data obtained using stable isotopes) is obviously lacking, an in-depth analysis also revealed several additional limitations: some values used were from individuals with bone disease; the impact of ethnicity was not discussed; the impact of the choice of calcium kinetic model (for experimental data analysis) was not considered; and finally, only calcium deposition data were considered and not the entire set of calcium-related data (deposition, release, net retention, and total mass).

Recently, we established a physiologically-based toxicokinetic model for aluminium built on a comprehensive animal and human toxicokinetic dataset [12, 13]. While expanding the model for use in regulatory risk assessment of aluminium exposure resulting from vaccines [14] and immunotherapeutics [15, 16], it became clear that dynamic changes in calcium kinetics during childhood become relevant due to the long half-life of several years for aluminium in bone [13]. The importance of accounting for age-related changes in physiologically-based models has also been described in a broader context [17].

The objective of this work was to expand the understanding on calcium kinetics in two ways: first, by reviewing literature data on total calcium mass, net calcium retention, calcium deposition and release rates from bone in males and females of different ages, including those not described in earlier studies. Second, by creating a consistent and ready-to-use equations describing calcium kinetics as a function of age and sex.

## 2. MATERIALS AND METHODS

### 2.1 Data collection and processing

To comprehensively retrieve data on calcium kinetic studies (deposition, release, and/or net retention), Google scholar was searched with the keywords “calcium kinetics” AND “stable isotope” AND “bone” AND “children” (April 2^nd^, 2024). The resulting 218 hits were screened for relevance. In addition, relevant cited and citing literature was screened.

The following inclusion criteria were applied:

- only healthy individuals (in particular, no bone diseases) aged under 40 years (i.e., no bone loss due to ageing),
- studies or study arms consisting mainly of white/Asian/hispanic individuals, since calcium kinetic rates have been shown to be increased in blacks [18, 19],
- sex reported (either individually or pooled),
- chronological age reported (not post-menarcheal age or age of peak growth velocity).

For each article, the number of study participants and their age range, sex and ethnicity was retrieved. If available, individual values on calcium mass or kinetics were extracted from tables or digitized from figures using WebPlotDigitizer [20]. If only summary statistics on age and calcium kinetics were reported, these were converted to mean ± SD assuming a normal distribution. Furthermore, study duration, calcium kinetic model and used isotopes used in the calcium kinetic study were recorded.

Data on total bone calcium mass were taken from two sources: (i) the ICRP publication 89 [21], which reported consolidated bone calcium masses for different ages (newborns, 1, 5, 10, 15 years, adults) and in males/females; and (ii) a comprehensive study by Ellis et al. [4], which reported whole-body bone mineral content (BMC) quantified by dual x-ray absorptiometry resolved yearly in age (4-18 years) and stratified by sex and ethnicity.

All calcium rates retrieved from literature, including the net retention rate 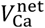, the deposition rate 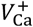 and the release rate 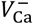, were converted (if necessary) to units kg Ca/year and mass measurements (*M*_Ca_) to units kg Ca. To convert BMC measurements to total calcium mass *M*_Ca_, we used the experimentally derived Ca fraction of BMC of 32.2% [3], which is lower than the theoretical Ca fraction of pure hydroxyapatite (39.15%) [22].

A schematic overview on experimental data and their link to the calcium kinetic rates is shown in Figure 1. Calcium deposition and release rates (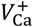 and 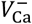) are not measured directly, but instead estimated from time-resolved calcium isotope ratios in blood, urine and/or stool via a calcium kinetic model (either compartmental or non-compartmental).

**Figure 1:**
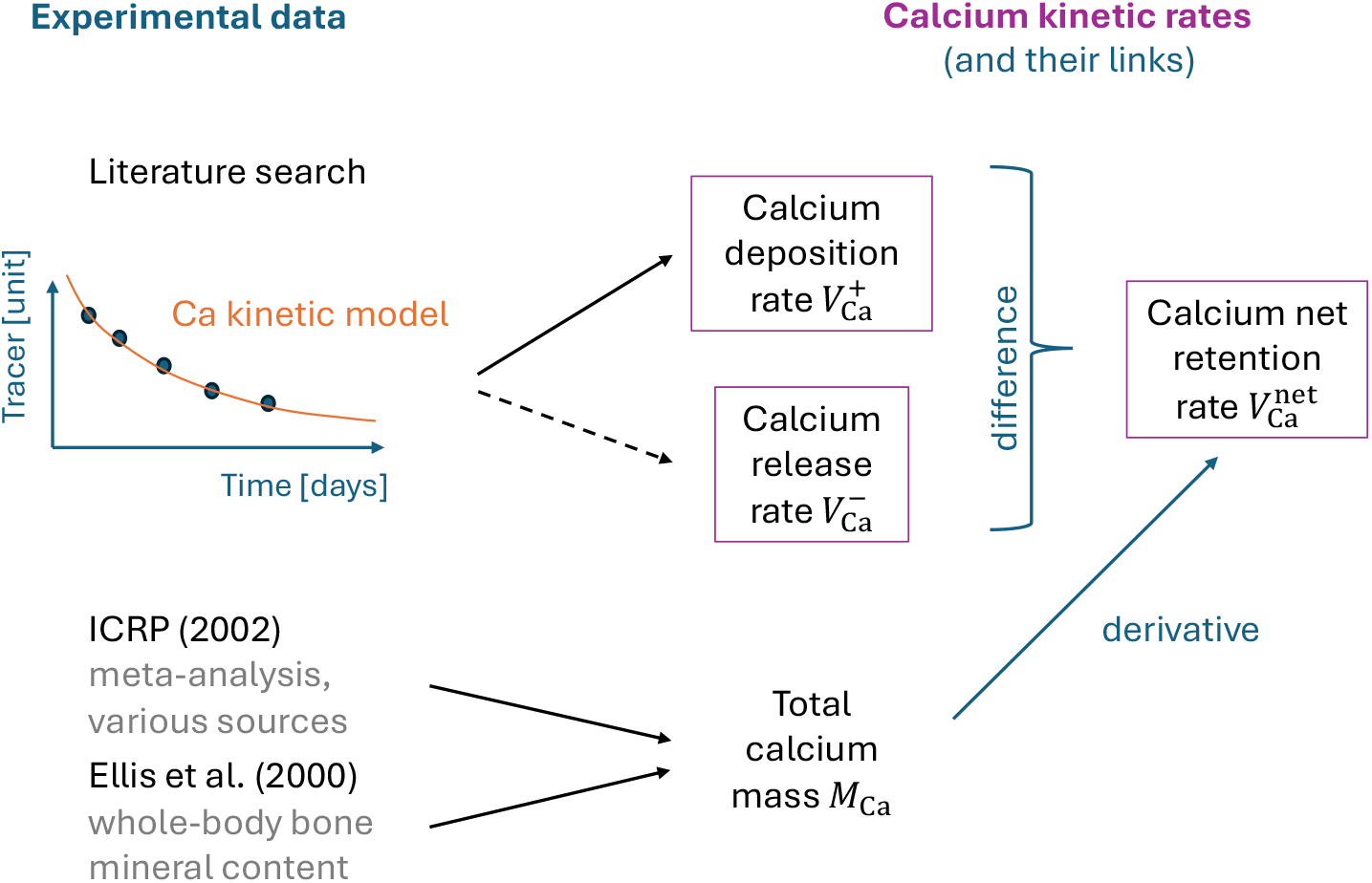
Overview on experimental data relating to calcium kinetics. Calcium tracer data are used to fit a calcium kinetic model, leading to an estimate for 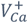 (and depending on the calcium model, possibly also 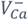). Total Ca mass data are taken from the ICRP report and computed from whole-body bone mineral content reported in Ellis et al. (2002). Calcium net retention is linked to both total calcium mass (as its derivative with respect to age) and calcium deposition and release (as their difference).

The impact of the choice of calcium kinetic model on estimated calcium rates has been discussed in the literature. While different compartmental models have been shown to generate similar kinetic estimates [23], the (non-compartmental) “sum of exponentials” model has been described to overfit the data by 15-25% [24, 25]. We therefore reduced all calcium kinetic rates obtained using this model by 20%.

### 2.2 Functions for calcium mass and kinetics

In order to build on the work of O’Flaherty in the 90s, total bone calcium mass *M*_Ca_ in kg Ca was predicted as a function of body weight (BW) depending on post-natal age *a* in years and sex [26]:

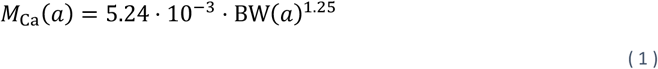

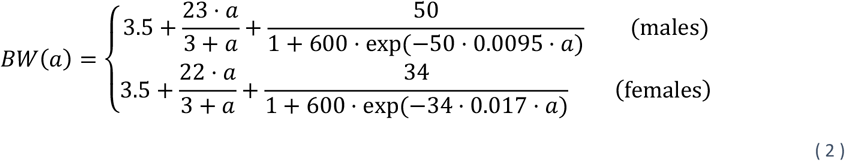

Subsequently, we investigated whether these functions are able to fit well to the ICRP data (used in many toxicokinetic models) and to other recent literature on total bone calcium mass [21, 4].

We then calculated net retention rate of bone calcium 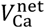 in kg Ca/year as the derivative of total bone calcium mass with respect to age,

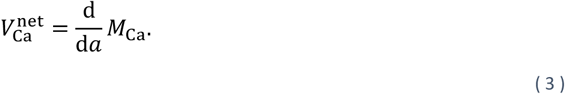

This prediction was then compared against the collected literature data.

Analogously to the approach used to describe calcium deposition in rats [27], a function for the calcium deposition rate 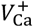 in kg Ca/year was established in two steps: First, data on (absolute) calcium deposition rate were converted to fractional calcium deposition rates 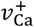 in 1/year via the relationship

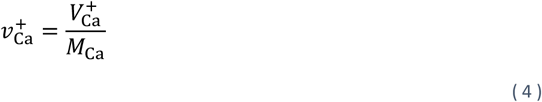

using the corresponding predicted total bone calcium mass *M*_Ca_ (i.e., same age and sex). Second, a functional relationship for fractional calcium deposition rate was developed based on the fractional calcium deposition rate data. While a sex-independent fractional rate was assumed for young children and for adults, sex-dependent rates were considered during puberty. The absolute calcium deposition rate 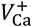 was then calculated from the fitted fractional calcium deposition rate, resulting in sex-independent rates for young children, but sex-specific rates thereafter (including adults) due to the sex-dependent total bone calcium mass *M*_Ca_. This two-step approach allowed to use simpler functional relationships and to more easily incorporate the assumption of sex-independent adult fractional calcium deposition rates, compared to a direct attempt to model 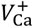 .

The calcium release rate from bone 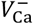 in kg Ca/year was determined as the difference between calcium deposition rate and calcium net retention rate,

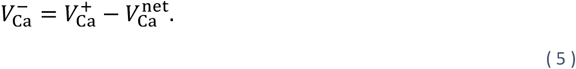

The resulting function was then compared against the collected literature data.

## 3. RESULTS

### 3.1 Data collection and processing

After a first screening of the literature, 19 references were identified that fulfilled the inclusion criteria. Of these, four were excluded after further investigation: one reference reported calcium deposition per kg body weight, but without providing individual body weight data [28]; another reference only contained information on preterm infants [29], which we considered out of scope for the current analysis; a third reference performed “longitudinal averaging”, i.e. averaging peak retention rates across different ages [30]; and in the fourth reference [19], the non-black subpopulation was identical to that of an earlier publication [31].

In two of the remaining 15 references, parts of the data were given as a function of post-menarcheal age, and these parts were therefore excluded [31, 25]. Another reference reported both individual rates in children, but without identifying them as boys or girls, as well as separate pooled rates for boys and girls [32]. In this case, the pooled data were used instead of the individual ones to allow for a sex-specific kinetic description. Furthermore, four references presented results on partly overlapping populations, but since different measures were reported (net retention and deposition rates derived from two different calcium models; partly pooled, partly individual data), all of them were kept in the dataset [18, 25, 33, 34].

All included references are summarized in Table 1. They cover the entire age range from newborns to adults, have slightly more data on females than males, are mostly conducted in white populations, contain only healthy individuals, and use a variety of setups for calcium kinetic studies. The collected/digitized data on Ca mass and rates are provided in tabular form as Supplementary Tables S1 and S2.

**Table 1:**
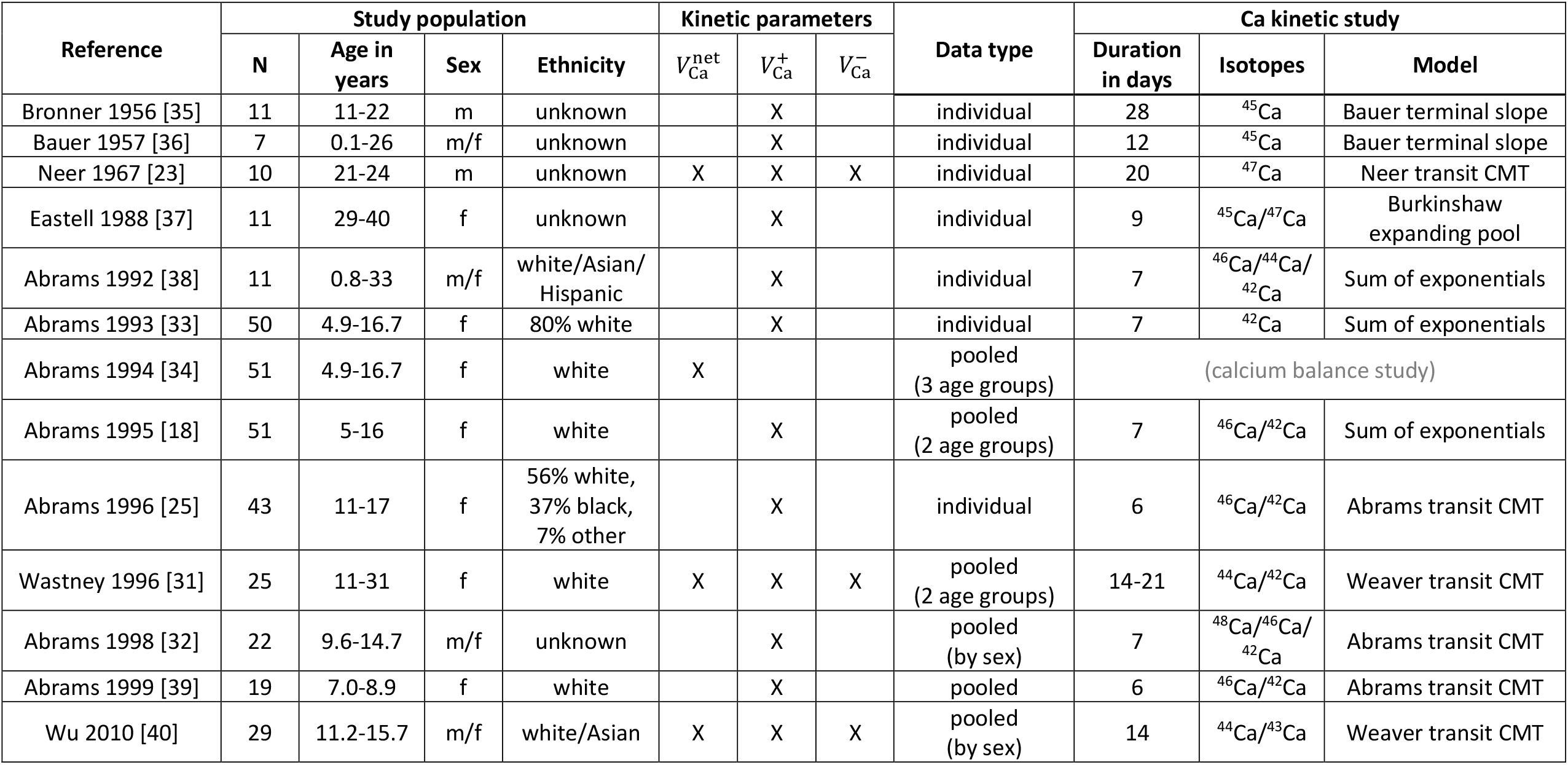
Summary of included references from the literature search. Abbreviations: m/f, male/female; CMT, compartment.

| Reference | Study population |  |  |  | Kinetic parameters |  |  | Data type | Ca kinetic study |  |  |
| --- | --- | --- | --- | --- | --- | --- | --- | --- | --- | --- | --- |
| | N | Age in years | Sex | Ethnicity | $V_{Ca}^{net}$ | $V_{Ca}^{+}$ | $V_{Ca}^{-}$ | | Duration in days | Isotopes | Model |
| Bronner 1956 [35] | 11 | 11-22 | m | unknown | | X | | individual | 28 | $^{45}Ca$ | Bauer terminal slope |
| Bauer 1957 [36] | 7 | 0.1-26 | m/f | unknown | | X | | individual | 12 | $^{45}Ca$ | Bauer terminal slope |
| Neer 1967 [23] | 10 | 21-24 | m | unknown | X | X | X | individual | 20 | $^{47}Ca$ | Neer transit CMT |
| Eastell 1988 [37] | 11 | 29-40 | f | unknown | | X | | individual | 9 | $^{45}Ca/^{47}Ca$ | Burkinshaw expanding pool |
| Abrams 1992 [38] | 11 | 0.8-33 | m/f | white/Asian/Hispanic | | X | | individual | 7 | $^{46}Ca/^{44}Ca/^{42}Ca$ | Sum of exponentials |
| Abrams 1993 [33] | 50 | 4.9-16.7 | f | 80% white | | X | | individual | 7 | $^{42}Ca$ | Sum of exponentials |
| Abrams 1994 [34] | 51 | 4.9-16.7 | f | white | X |  |  | pooled (3 age groups) | (calcium balance study) |  |  |
| Abrams 1995 [18] | 51 | 5-16 | f | white | | X | | pooled (2 age groups) | 7 | $^{46}Ca/^{42}Ca$ | Sum of exponentials |
| Abrams 1996 [25] | 43 | 11-17 | f | 56% white, 37% black, 7% other | | X | | individual | 6 | $^{46}Ca/^{42}Ca$ | Abrams transit CMT |
| Wastney 1996 [31] | 25 | 11-31 | f | white | X | X | X | pooled (2 age groups) | 14-21 | $^{44}Ca/^{42}Ca$ | Weaver transit CMT |
| Abrams 1998 [32] | 22 | 9.6-14.7 | m/f | unknown | | X | | pooled (by sex) | 7 | $^{48}Ca/^{46}Ca/^{42}Ca$ | Abrams transit CMT |
| Abrams 1999 [39] | 19 | 7.0-8.9 | f | white | | X | | pooled | 6 | $^{46}Ca/^{42}Ca$ | Abrams transit CMT |
| Wu 2010 [40] | 29 | 11.2-15.7 | m/f | white/Asian | X | X | X | pooled (by sex) | 14 | $^{44}Ca/^{43}Ca$ | Weaver transit CMT |

### 3.3 Functional description of calcium data

Data on total bone calcium mass, as well as predictions based on the O’Flaherty calcium mass function (1)-(2), are depicted in Figure 2. Predictions were found to be in good agreement with the data, for both boys and girls, from birth to adulthood. Therefore, no further adjustment was done to this part of the calcium functional description.

**Figure 2:**
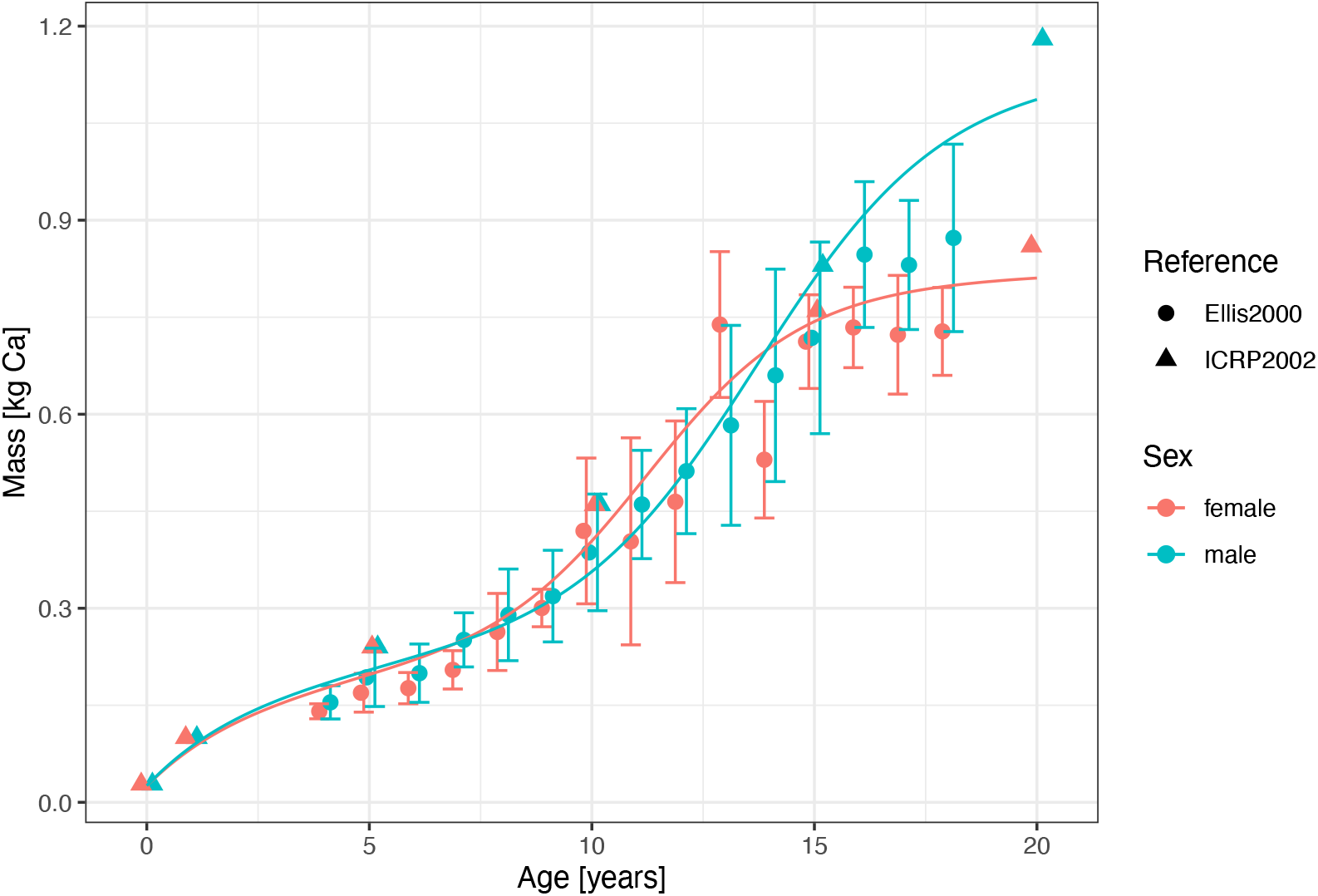
Comparison of predictions using the calcium mass function 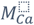 calculated via (1)-(2) to data on total calcium mass, as a function of post-natal age. For visualization, the ICRP “adult” age is set to 20 years.

The derived calcium net retention function based on (*3*) is shown in Figure 3, together with independent calcium net retention data. Variability in the data is large, but the overall trend is captured well in the calcium net retention function. Although peak values in Wu (2010) seem larger than predicted (no estimate of variability is reported though in Wu (2010)), the difference between boys and girls is represented well.

**Figure 3:**
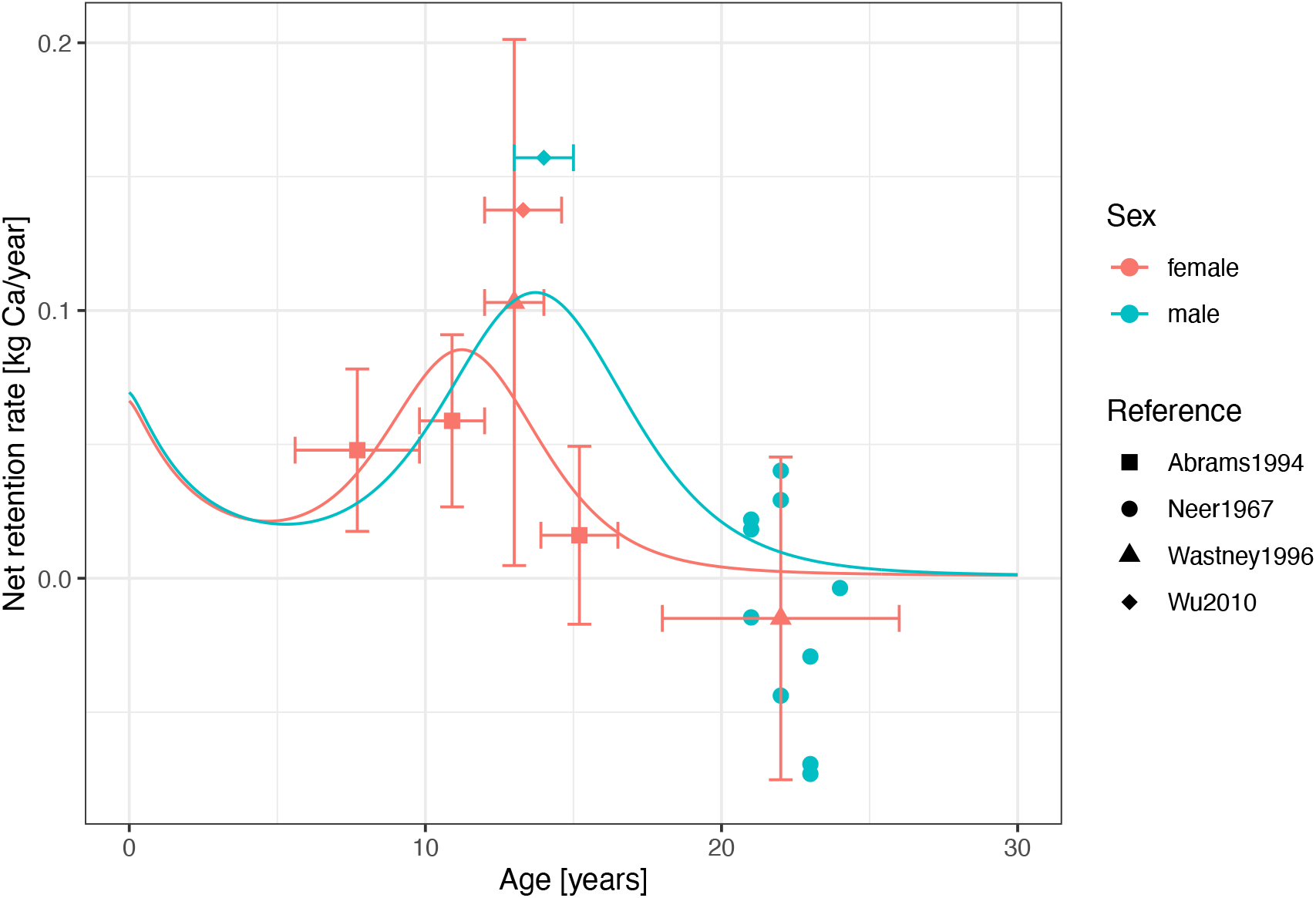
Comparison of predictions using the calcium net retention rate function 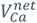 calculated via (3) to data on calcium net retention, as a function of post-natal age.

Subsequently, the large data collection of calcium deposition rates obtained via kinetic studies was described. As stated in the Methods section, a function was first developed for fractional calcium deposition rate 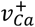 in 1/year. Different shapes of fractional deposition curves 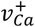 were tested, and the final curve (shown in Supplementary Figure S1) is given by the following equation (with post-natal age *a* in years and 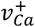 in 1/year):

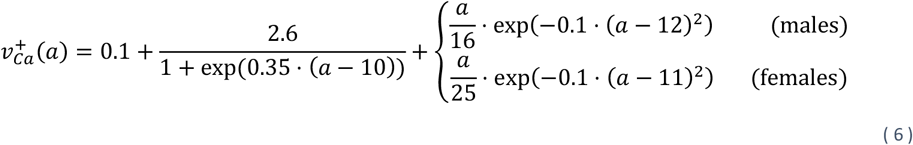

It uses an adult baseline of 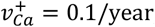, a sex-independent decreasing term describing rapid fractional calcium deposition during early childhood (starting from 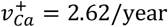 in newborns), and a sex-specific acceleration during puberty at the age of 12 years in boys and 11 years in girls.

The absolute calcium deposition rate 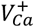 in kg Ca/year was obtained subsequently by solving (4) for 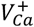; the resulting curve is shown in Figure 4. The overall trend in the calcium deposition data 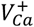 is described well with this function. Compared to the previously published calcium deposition function [11], the peak at puberty is halved and the difference between boys and girls is more pronounced. The main reasons for these descrepancies are (i) the relationship in [11] was build on deposition rates that originated from the “sum of exponentials” calcium kinetic model and were thus too high; (ii) the removal of large deposition rates from two black children in our dataset; and (iii) the inclusion of newer data describing differences between boys and girls in our analysis. After reducing all deposition data using the “sum of exponentials” model by 20% as described in the methods, no systematic differences between different calcium kinetic models were seen.

**Figure 4:**
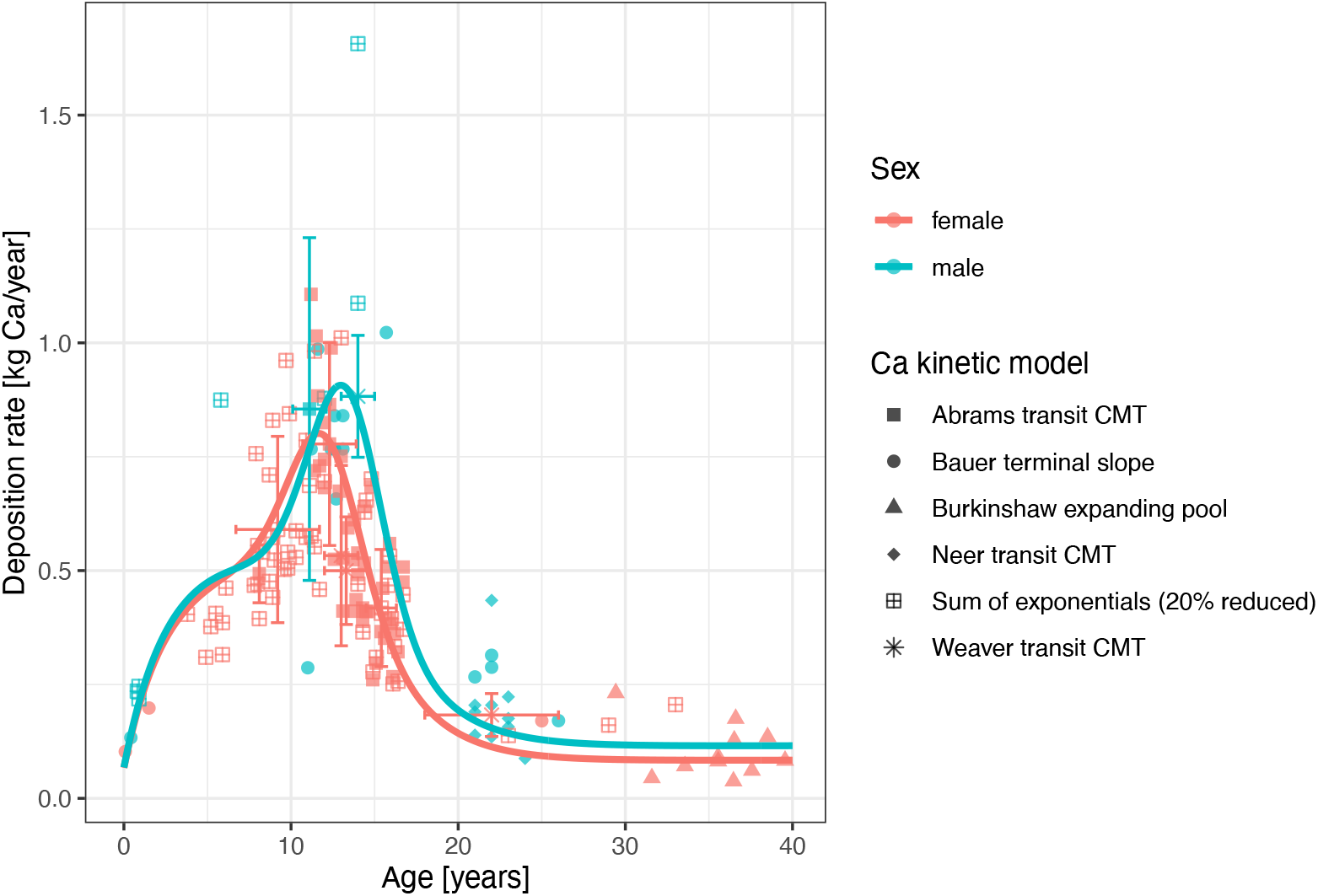
Comparison of predictions using the calcium deposition rate function 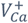, obtained by solving (4) for 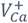, to data on calcium deposition, as a function of post-natal age. Due to the large number of references included, calcium kinetic models rather than references themselves are shown in the legend (full details including a reference for each data point are given as supplementary material).

The calcium release rate from bone 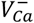, calculated via (5), was then compared with literature data (Figure 5). Again, the overall trend was captured well, indicating that the different calcium functions (total mass, net retention/deposition/release rates) are consistent in their description of the calcium datasets.

**Figure 5:**
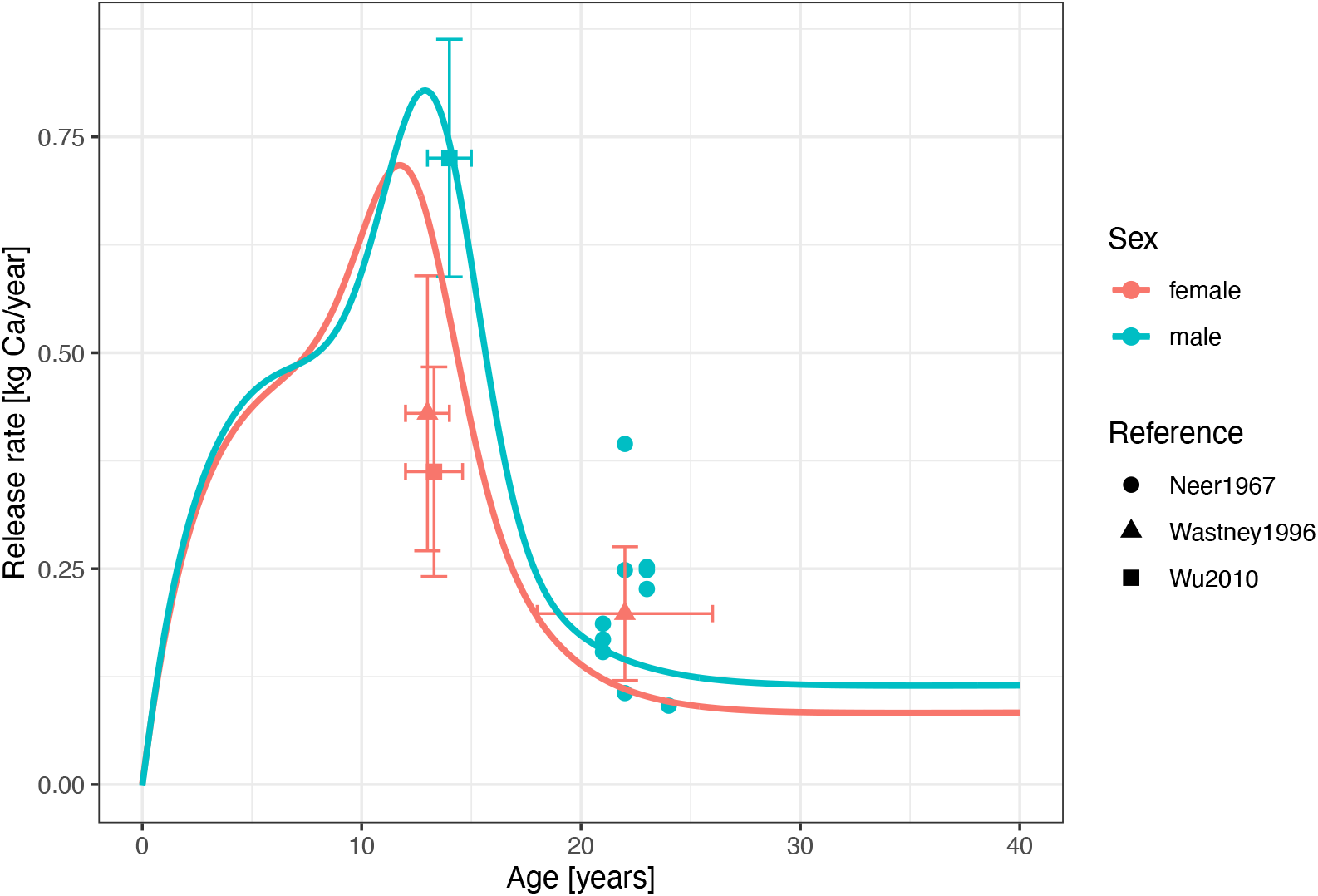
Comparison of predictions using the calcium release rate function 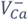 calculated via (5) to data on calcium release, as a function of post-natal age.

## 4. DISCUSSION

We have compiled a comprehensive dataset on calcium kinetics in human bone, from mainly Caucasian healthy term newborns to adults and in males and females. In the dataset, consistent age- and sex-specific changes can be seen over a variety of studies and analysis methods. Describing these changes using functional relationships, we provided a consolidated mathematical description of calcium kinetics, which is expected to be a valuable tool in toxicokinetic analyses of bone-seeking elements. A description of calcium kinetics as a continuous function of age also allowed to successfully link these data to total calcium mass, which is important when incorporating calcium kinetic rates into physiologically-based models. In addition, the compiled dataset on calcium kinetics may form a basis for further analyses, for example including the impact of diseases, ageing or nutritional research.

The large heterogeneity of the collected data also allowed to investigate the impact of different population and study characteristics. The duration of kinetic studies differed between laboratories, with some conducting one-week and others two-week studies. While the validity of deposition rate estimates for 7-day kinetic studies has been questioned [41, 42], no such trend could be seen in the compiled dataset. Furthermore, reducing by 20% all deposition rates obtained via the sum of exponentials model resulted a good alignment of deposition data obtained via different kinetic models, without any systematic outlier (see Figure 4).

The functions for calcium kinetics have been built from data in mainly Caucasian populations. No major differences are expected in Asian (similar estimates in whites [31] and Asians [40]) or Hispanic populations (no significant difference found in [39]). This contrasts with the calcium kinetics in blacks, where all rates are considerably larger [18, 19]. Unfortunately, the literature on calcium kinetics in blacks is too sparse to formulate specific functions in the same way.

While a central age- and sex-dependent trend for calcium kinetics can be clearly identified in the calcium kinetics dataset, there is considerable between- and within-study variability. Most likely, differences in oral bioavailability of calcium contributed significantly to this variability. Calcium absorption has not been investigated in the current work, and coupling the calcium kinetic functions to a calcium absorption function would be an important extension with relevance for nutrition sciences. Another major source of variability is differences in body weight (for the same age and sex). By normalizing data to sex- and age-specific reference body weights, this effect could be compensated for. However, many references did not report individual body weights alongside the calcium kinetic rates and hence, such an analysis was not conducted here.

For older adults, especially post-menopausal women, bone resorption increases while bone formation stays approximately constant, thus leading to net bone loss [43, 44]. The current functional descriptions of calcium kinetics do not account for this effect and hence, should not be used outside the validated age range of 0 to 40 years without a critical discussion. Similarly, an extension would be required for preterm infants.

## Funding

This research did not receive any specific grant from funding agencies in the public, commercial, or not-for-profit sectors.

## Supplementary Material

Supplementary Table S1: Collection of literature data on calcium mass.

Supplementary Table S2: Collection of literature data on calcium net retention, deposition, and release rates.

Supplementary Figure S1: Fit to fractional calcium deposition rates

**Supplementary Figure S1:**
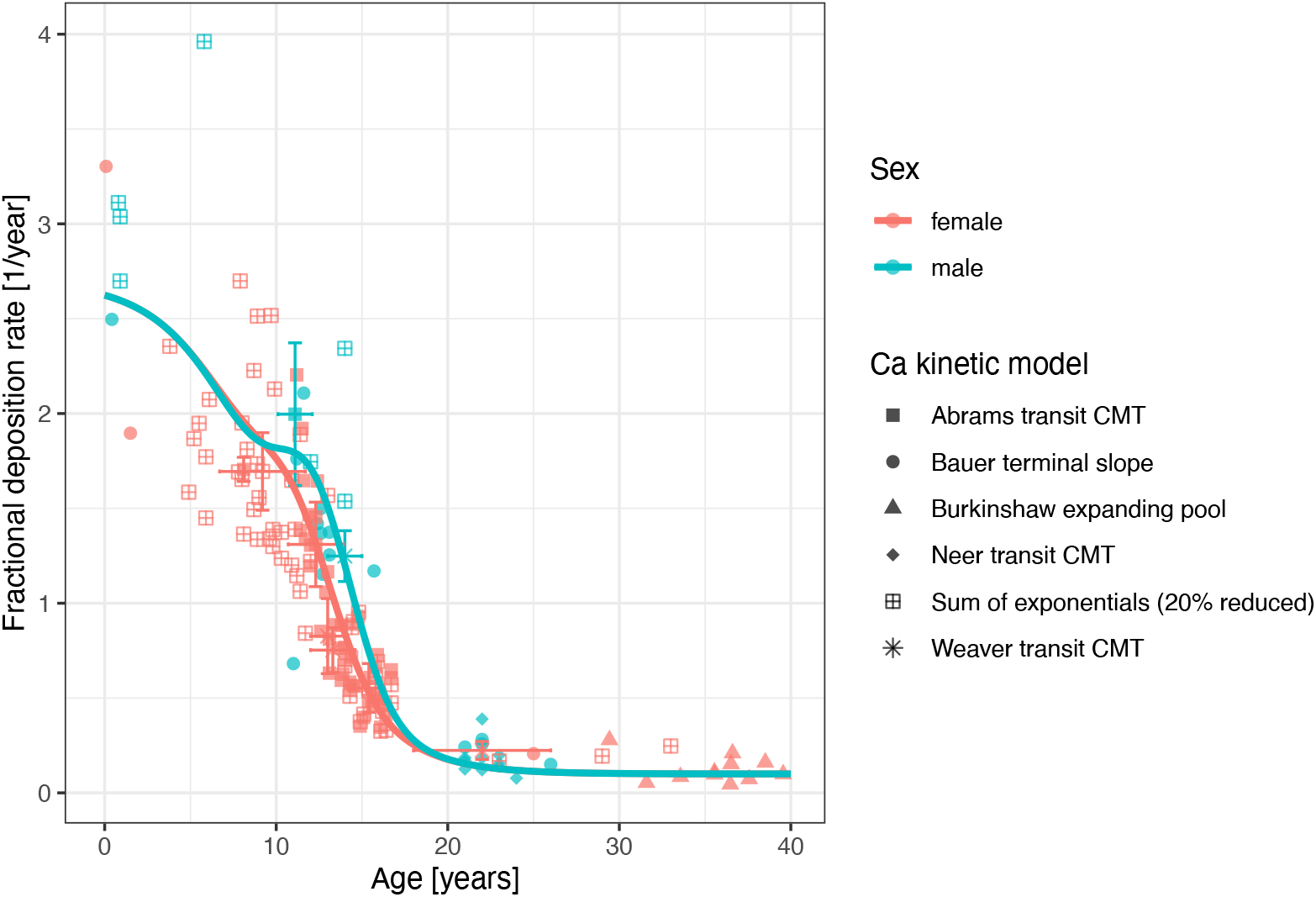
Comparison of predictions using the fractional calcium deposition rate function 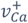 calculated via (6) to the (transformed) data on fractional calcium deposition, as a function of post-natal age.

## Notes

### Competing Interest Statement

The authors have declared no competing interest.

## References

[1] S. A. Abrams, “Calcium turnover and nutrition through the life cycle,” Proceedings of the Nutrition Society, vol. 60, pp. 283–289, 2001.

[2] L. S. Hillman, E. Tack, D. G. Covell, N. E. Vieira and A. L. Yergey, “Measurement of True Calcium Absorption in Premature Infants Using Intravenous 46Ca and Oral 44Ca,” Pediatric Research, vol. 23, pp. 589–594, 1988.

[3] K. J. Ellis, R. J. Shypailo, A. Hergenroeder, M. Perez and S. Abrams, “Total body calcium and bone mineral content: Comparison of dual-energy X-ray absorptiometry with neutron activation analysis,” Journal of Bone and Mineral Research, vol. 11, p. 843–848, June 1996.

[4] K. J. Ellis, R. J. Shypailo, S. A. Abrams and W. W. Wong, “The Reference Child and Adolescent Models of Body Composition: A Contemporary Comparison,” Annals of the New York Academy of Sciences, vol. 904, p. 374–382, May 2000.

[5] C. Weaver, A. Rothwell and K. Wood, “Measuring calcium absorption and utilization in humans,” Current Opinion in Clinical Nutrition and Metabolic Care, vol. 9, p. 568–574, 2006.

[6] E. J. O’Flaherty, “A physiologically based kinetic model for lead in children and adults.,” Environmental Health Perspectives, vol. 106, p. 1495–1503, December 1998.

[7] F. Bronner, “Bone and calcium homeostasis,” Neurotoxicology, pp. 13(4):775–782, 1992.

[8] R. B. Richardson, “A physiological skeletal model for radionuclide and stable element biokinetics in children and adults,” Health Physics, vol. 99, p. 471–482, October 2010.

[9] R. W. Leggett, “A Generic Age-Specific Biokinetic Model for Calcium-like Elements,” Radiation Protection Dosimetry, vol. 41, p. 183–198, June 1992.

[10] H. Pertinez, M. Chenel and L. Aarons, “A Physiologically Based Pharmacokinetic Model for Strontium Exposure in Rat,” Pharmaceutical Research, vol. 30, pp. 1536–1552, 2013.

[11] E. J. O’Flaherty, “Physiologically Based Models of Metal Kinetics,” Critical Reviews in Toxicology, vol. 28, p. 271–317, January 1998.

[12] K. Weisser, S. Stübler, W. Matheis and W. Huisinga, “Towards toxicokinetic modelling of aluminium exposure from adjuvants in medicinal products,” Regulatory Toxicology and Pharmacology, vol. 88, p. 310–321, August 2017.

[13] C. Hethey, N. Hartung, G. Wangorsch, K. Weisser and W. Huisinga, “Physiology-based toxicokinetic modelling of aluminium in rat and man,” Archives of Toxicology, vol. 95, p. 2977–3000, August 2021.

[14] K. Weisser, T. Göen, J. Oduro, G. Wangorsch, K. Hanschmann and B. Keller-Stanislawski, “Aluminium in plasma and tissues after intramuscular injection of adjuvanted human vaccines in rats.,” Archives of Toxicology, vol. 93, pp. 2787–2796, 2019.

[15] K. Weisser, T. Göen, J. Oduro, G. Wangorsch, H. K.M.O., B. and Keller-Stanislawski, “Aluminium from adjuvanted subcutaneous allergen immunotherapeutics is mainly detected in bone,” Allergy, vol. 75, pp. 215–217, 2020.

[16] K. Weisser, “Toxicokinetics of aluminium - novel insights in an old adjuvant,” Allergo Journal International, 2024.

[17] K. Price, S. Haddad and K. Krishnan, “Physiological modeling of age-specific changes in the pharmacokinetics of organic chemicals in children.,” J Toxicol Environ Health A, vol. 66, pp. 417–433, 2003.

[18] S. A. Abrams, K. O. O’Brien, L. K. Liang and J. E. Stuff, “Differences in calcium absorption and kinetics between black and white girls aged 5–16 years,” Journal of Bone and Mineral Research, vol. 10, p. 829–833, May 1995.

[19] R. J. Bryant, M. E. Wastney, B. R. Martin, O. Wood, G. P. McCabe, M. Morshidi, D. L. Smith, M. Peacock and C. M. Weaver, “Racial Differences in Bone Turnover and Calcium Metabolism in Adolescent Females,” The Journal of Clinical Endocrinology & Metabolism, vol. 88, p. 1043–1047, March 2003.

[20] A. Rohatgi, “WebPlotDigitizer,” 2022. [Online]. Available: https://automeris.io/WebPlotDigitizer.

[21] ICRP, “Basic anatomical and physiological data for use in radiological protection: reference values. ICRP Publication 89,” Annals of the ICRP, vol. 32, no. 3-4, p. 1–277, September 2002.

[22] M. Markovic, B. O. Fowler and M. S. Tung, “Preparation and comprehensive characterization of a calcium hydroxyapatite reference material,” Journal of Research of the National Institute of Standards and Technology, vol. 109, p. 553, November 2004.

[23] R. Neer, M. Berman, L. Fisher and L. E. Rosenberg, “Multicompartmental Analysis of Calcium Kinetics in Normal Adult Males,” Journal of Clinical Investigation, vol. 46, p. 1364–1379, August 1967.

[24] J.-P. Aubert, F. Bronner and L. J. Richelle, “Quantitation of calcium metabolism. Theory,” Journal of Clinical Investigation, vol. 42, p. 885–897, June 1963.

[25] S. A. Abrams, K. O. O’Brien and J. E. Stuff, “Changes in calcium kinetics associated with menarche.,” The Journal of Clinical Endocrinology & Metabolism, vol. 81, p. 2017–2020, June 1996.

[26] E. J. O’Flaherty, “Physiologically based models for bone-seeking elements: III. Human skeletal and bone growth,” Toxicology and Applied Pharmacology, vol. 111, p. 332–341, November 1991.

[27] E. J. O’Flaherty, “Physiologically based models for bone-seeking elements: II. Kinetics of lead disposition in rats,” Toxicology and Applied Pharmacology, vol. 111, p. 313–331, November 1991.

[28] R. P. Heaney and G. D. Whedon, “Radiocalcium studies of bone formation rate in human metabolic bone disease,” The Journal of Clinical Endocrinology & Metabolism, vol. 18, p. 1246–1267, November 1958.

[29] S. A. Abrams, R. J. Schanler, A. L. Yergey, N. E. Vieira and F. Bronner, “Compartmental Analysis of Calcium Metabolism in Very-Low-Birth-Weight Infants,” Pediatric Research, vol. 36, p. 424–428, October 1994.

[30] D. A. Bailey, A. D. Martin, H. A. McKay, S. Whiting and R. Mirwald, “Calcium Accretion in Girls and Boys During Puberty: A Longitudinal Analysis,” Journal of Bone and Mineral Research, vol. 15, p. 2245–2250, November 2000.

[31] M. E. Wastney, J. Ng, D. Smith, B. R. Martin, M. Peacock and C. M. Weaver, “Differences in calcium kinetics between adolescent girls and young women,” American Journal of Physiology-Regulatory, Integrative and Comparative Physiology, vol. 271, p. R208–R216, July 1996.

[32] S. A. Abrams, “The Relationship Between Magnesium and Calcium Kinetics in 9-to 14-Year-Old Children,” Journal of Bone and Mineral Research, vol. 13, p. 149–153, January 1998.

[33] S. A. Abrams, “Pubertal Changes in Calcium Kinetics in Girls Assessed Using 42Ca,” Pediatric Research, vol. 34, p. 455–459, October 1993.

[34] S. A. Abrams and J. E. Stuff, “Calcium metabolism in girls: current dietary intakes lead to low rates of calcium absorption and retention during puberty,” The American Journal of Clinical Nutrition, vol. 60, p. 739–743, November 1994.

[35] F. Bronner and R. S. Harris, “Absorption and metabolism of calcium in human beings, studied with calcium-45,” Annals of the New York Academy of Sciences, vol. 64, p. 314–325, August 1956.

[36] G. C. H. Bauer, A. Carlsson and B. Lindquist, “Bone Salt Metabolism in Humans Studied by Means of Radiocalcium,” Acta Medica Scandinavica, vol. 158, p. 143–150, January 1957.

[37] R. Eastell, P. D. Delmas, S. F. Hodgson, E. F. Eriksen, K. G. Mann and B. L. Riggs, “Bone Formation Rate in Older Normal Women: Concurrent Assessment with Bone Histomorphometry, Calcium Kinetics, and Biochemical Markers,” The Journal of Clinical Endocrinology & Metabolism, vol. 67, p. 741–748, October 1988.

[38] S. A. Abrams, N. V. Esteban, N. E. Vieira, J. B. Sidbury, B. L. Specker and A. L. Yergey, “Developmental changes in calcium kinetics in children assessed using stable isotopes,” Journal of Bone and Mineral Research, vol. 7, p. 287–293, March 1992.

[39] S. A. Abrams, K. C. Copeland, S. K. Gunn, J. E. Stuff, L. L. Clarke and K. J. Ellis, “Calcium Absorption and Kinetics Are Similar in 7- and 8-Year-Old Mexican-American and Caucasian Girls Despite Hormonal Differences,” The Journal of Nutrition, vol. 129, p. 666–671, March 1999.

[40] L. Wu, B. R. Martin, M. M. Braun, M. E. Wastney, G. P. McCabe, L. D. McCabe, L. A. DiMeglio, M. Peacock and C. M. Weaver, “Calcium requirements and metabolism in Chinese-American boys and girls,” Journal of Bone and Mineral Research, vol. 25, p. 1842–1849, July 2010.

[41] C. M. Weaver, M. Wastney and L. A. Spence, “Quantitative Clinical Nutrition Approaches to the Study of Calcium and Bone Metabolism,” in Nutrition and Bone Health, Springer New York, 2014, p. 361–377.

[42] C. M. Weaver, “Use of Calcium Tracers and Biomarkers to Determine Calcium Kinetics and Bone Turnover,” Bone, vol. 22, p. 103S–104S, May 1998.

[43] E. J. O’Flaherty, “Modeling Normal Aging Bone Loss, with Consideration of Bone Loss in Osteoporosis,” Toxicological Sciences, vol. 55, no. 1, pp. 171–188, 2000.

[44] L. A. Spence, E. R. Lipscomb, J. Cadogan, B. Martin, M. E. Wastney, M. Peacock and C. M. Weaver, “The effect of soy protein and soy isoflavones on calcium metabolism in postmenopausal women: a randomized crossover study,” The American Journal of Clinical Nutrition, vol. 81, p. 916–922, April 2005.

[45] E. J. O’Flaherty, “Physiologically based models for bone-seeking elements: IV. Kinetics of Lead Disposition in Humans,” Toxicology and Applied Pharmacology, vol. 118, p. 16–29, January 1993.

[46] F. Bronner, L. J. Richelle, P. D. Saville, J. A. Nicholas and J. R. Cobb, “Quantitation of calcium metabolism in postmenopausal osteoporosis and in scoliosis,” Journal of Clinical Investigation, vol. 42, p. 898–905, June 1963.

